# Strain-resolved CRISPRi-Seq reveals conserved antibiotic vulnerabilities in *Staphylococcus aureus*

**DOI:** 10.1101/2025.10.13.682098

**Authors:** Maria-Vittoria Mazzuoli, Jessica Burnier, Albane Schmid, Louise Martin, Absalom Janssen, Clement Gallay, Annelies S Zinkernagel, Jan-Willem Veening

**Author notes:** Correspondence to Maria Vittoria Mazzuoli or Jan-Willem Veening. These authors contributed equally (Maria-Vittoria Mazzuoli, Jessica Burnier).

## Abstract

*Staphylococcus aureus* remains a major clinical threat due to rising antibiotic resistance and high rates of treatment failure. Deciphering the genetic responses to antibiotic pressure and identifying conserved vulnerabilities are essential steps toward developing broadly effective therapies. Here, we constructed strain-resolved CRISPR interference (CRISPRi) libraries targeting all genes in four clinically relevant *S. aureus* strains spanning major clonal complexes. CRISPRi-seq screens enabled high-resolution mapping of their fitness landscapes and the definition of a core essentialome representing robust targets for antimicrobial intervention. Exposure of the CRISPRi libraries to four mechanistically distinct antibiotics revealed genome-wide susceptibility profiles, identifying both strain-dependent and conserved susceptibility signatures shaped by the drug mode of action and genetic background. Analysis of these conserved vulnerabilities provided insight into antibiotic-specific stress responses and resistance mechanisms. Among the core determinants of vancomycin vulnerability, we identified several previously uncharacterized genes, including a conserved membrane-associated operon, here designated EsrABC, whose disruption markedly increases vancomycin sensitivity in the four strains. Our study provides a genome-wide atlas of *S. aureus* fitness and conditional vulnerabilities, fully explorable in the here-developed online AureoBrowse platform (https://aureobrowse.veeninglab.com/), revealing candidates for synergistic therapies and potential therapeutic targets.

## Introduction

The global spread of antibiotic resistance poses a growing threat to public health, exacerbated by a declining pipeline of new antibiotics and vaccines. Among resistant pathogens, *Staphylococcus aureus* is of particular concern due to its high resistance rates, virulence, and global burden of disease ^1–3^. Methicillin-resistant *S. aureus* (MRSA) is a leading cause of hospital- and community-acquired infections, including pneumonia, sepsis, endocarditis, and prosthetic device infections. Vancomycin remains the first-line treatment for MRSA, but the emergence of vancomycin-intermediate (VISA) and resistant (VRSA) strains ^4–6^ underscores the need to better understand resistance mechanisms and identify alternative therapeutic strategies. Broadly effective therapies against *S. aureus* require targets that are conserved across strains. Yet, gene essentiality is highly context-dependent, varying with genetic background and environmental conditions ^7^. Defining essential pathways in diverse *S. aureus* lineages is therefore critical for identifying robust drug targets. Targeting genes that become conditionally essential under antibiotic pressure offers an additional strategy to design synergistic drug combinations that potentiate antibacterial activity^8–10^.

Functional genomics, through systematic mutagenesis and genome-wide screening, provides a powerful approach to uncover essential and conditionally essential genes. When combined with antibiotic stress, chemogenomic approaches enable profiling of drug-specific responses and the identification of potential targets for combination therapies ^11^. Transposon sequencing (Tn-seq) has been widely used in bacteria to identify essential genes and susceptibility determinants^7,12–15^. However, because mutants lacking essential genes are underrepresented in transposon libraries, Tn-seq provides limited resolution for studying core processes targeted by antibiotics. CRISPR interference sequencing (CRISPRi-seq) addresses this limitation by allowing tunable, conditional repression of gene expression, enabling the analysis of essential genes. CRISPRi relies on the co-expression of a catalytically inactive Cas9 protein (dCas9) and a single-guide RNA (sgRNA) that directs dCas9 to the target locus to block transcription without DNA cleavage^16^. Previous studies have applied CRISPRi-seq to identify antibiotic susceptibility determinants across bacterial species and drugs, including dalbavancin and amoxicillin in *S. aureus* ^17^. However, existing *S. aureus* CRISPRi libraries typically cover only a subset of genes, are based on single laboratory strains, or are designed at the operon level. These limitations restrict genome-wide, gene-level comparisons across genetically diverse clinical isolates, particularly given incomplete operon annotations.

In this study, we developed strain-specific CRISPRi libraries for four *S. aureus* isolates encompassing distinct clonal complexes and virulence capacities: NCTC8325-4 (CC8), CL-2136 (CC45), JE2 (CC8), and Cowan I (CC30). NCTC8325-4 is a prophage-cured derivative of the widely used laboratory strain NCTC8325 ^18^, whereas CL-2136 was recently isolated from a bloodstream infection at the University Hospital of Zurich, Switzerland. JE2 is a plasmid-cured derivative of the community-acquired MRSA strain USA300, a virulent and fluoroquinolone-resistant isolate, whereas Cowan I is a methicillin-sensitive *S. aureus* strain with defective *agr* signaling and attenuated virulence^19,20^. CRISPRi-seq screening across these isolates enabled high-resolution mapping of fitness determinants, expanding the known *S. aureus* fitness landscape and revealing extensive strain-specific gene dependencies. A genome-wide atlas of core and strain-dependent essentialomes is available through an interactive browser (https://aureobrowse.veeninglab.com). We used this platform to examine gene requirements under antibiotic stress in MRSA JE2 and MSSA Cowan I. Four clinically relevant antibiotics with distinct mechanisms of action were selected: flucloxacillin (β-lactam), vancomycin (glycopeptide), levofloxacin (fluoroquinolone), and rifampicin (rifamycin), targeting, respectively, cell wall synthesis, DNA replication, and transcription^21–24^. All four antibiotics are commonly used in *S. aureus* infections. Levofloxacin and rifampicin are potent against biofilm-associated infections and are typically combined since they rapidly select for resistance when used alone^25,26^. These features underscore the importance of evaluating their potential in combination therapies.

Here, we optimized the first genome-wide, high-coverage, strain-resolved CRISPRi-seq platform and applied it to four diverse *S. aureus* strains, including a recently isolated clinical strain. CRISPRi-seq screenings across these genetic backgrounds enabled us to define a core essentialome, representing promising targets for broadly effective antibacterial strategies. In addition, this work highlighted the strain-dependent nature of fitness landscapes. CRISPRi-seq performed in two divergent strains (Cowan and JE2) exposed to four mechanistically distinct antibiotics (flucloxacillin, vancomycin, levofloxacin and rifampicin) revealed both strain-specific and conserved genome-wide susceptibility profiles, shaped by both drug mechanism and genetic background. Analysis of these conserved vulnerabilities provided insight into antibiotic-specific stress responses and resistance mechanisms. Among the core determinants of vancomycin susceptibility, we identified multiple previously uncharacterized genes, including a conserved membrane-associated operon, here designated EsrABC, whose disruption markedly increases vancomycin sensitivity in the four strains. Together, these findings provide a framework for developing new strategies to combat the growing threat of vancomycin resistance in *S. aureus*.

## Results

### A switchable, chromosome-encoded CRISPRi system to explore gene requirements

In our previous work, we developed a two-plasmid, IPTG-inducible CRISPRi system and used it to perform CRISPRi-seq in *S. aureus* strain NCTC8325-4 ^17^. More recently, it was shown that reducing the copy number of *dcas9* to a single chromosomal copy by integrating it into the *spa* locus resulted in tighter and more tunable expression control^27^. To further optimize the system, we integrated a markerless *dcas9* variant into a newly designated intergenic *sep* locus (Staphylococcal Expression Platform; see Methods) of the *S. aureus* genome. This variant is controlled by a tetracycline-inducible promoter responsive to anhydrotetracycline (aTc) and doxycycline (Fig. 1A). Using a pMAD-based plasmid, we inserted the *Ptet-dcas9* construct into the chromosomes of four *S. aureus* strains: NCTC8325-4, JE2, Cowan I, and CL-2136. Expression of *dcas9* had no detectable impact on growth across these genetic backgrounds (Supplementary Fig. 1A). For sgRNA delivery, we modified the previously described pVL2336 plasmid ^17^ by introducing a tetracycline resistance cassette (*tetO*), generating the vector pVL4930 (Fig. 1B). This ensures that growth of sgRNA clones are not negatively affected by the TetR inducer. To assess the system’s ability to knock-down a gene required for growth, we targeted *hu*, which encodes the essential conserved histone-like protein HU, in the four strains. A control sgRNA containing a random sequence served as a negative control. As shown in Fig. 1C, increasing aTc concentrations induced *dcas9* expression, leading to pronounced growth inhibition in strains carrying *hu*-targeting sgRNAs, but not in those harboring the control sgRNA. These results establish a functional, tightly regulated, and inducible gene knockdown system suitable for both laboratory and clinical *S. aureus* strains.

**Figure 1.**
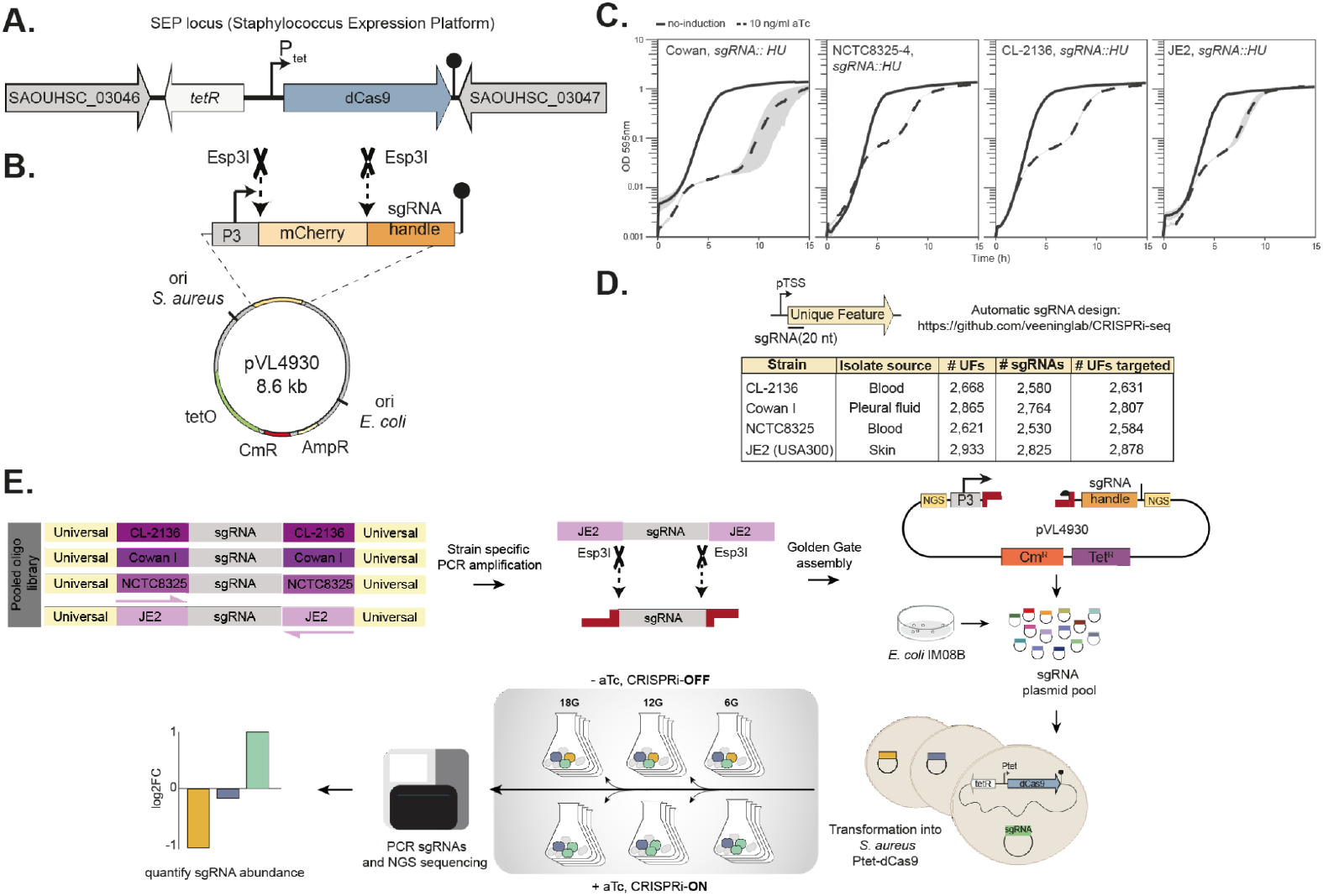
Construction of a CRISPRi-optimized system for clinically relevant *S. aureus* strains. Schematic representation of the *sep* locus harbouring the *sep::Ptet-dcas9* construct. **B)** Map of the sgRNA cloning vector pVL4930. **C)** CRISPRi-mediated knockdown of an essential gene (*hu*) in *S. aureus* strains NCTC8325-4, JE2, Cowan I, and CL-2136 resulted in pronounced growth defects. **D)** Overview of the sgRNA design strategy enabling genome-wide, strain-specific conditional gene knockdown. **E)** Outline of the workflow for constructing high-throughput, synthetically pooled sgRNA libraries. Strain-specific sgRNAs were amplified and inserted into pVL4930; the flanking NGS boxes represent Illumina adapter sequences used for sequencing. Resulting plasmids were first transformed into a methylation-permissive *E. coli* strain to bypass *S. aureus* restriction barriers and subsequently introduced into *S. aureus* strains carrying the *sep::Ptet-dcas9* construct. CRISPRi-seq screenings were then performed over 6, 12 and 18 generations in the presence or absence of the inducer (anhydrotetracycline or doxycycline). Plasmids were recovered, and sgRNA sequences were amplified with Illumina adapters, sequenced, and quantified.

### Strain-specific libraries capture the diversity of clinical *S. aureus*

We next designed strain-specific pooled CRISPRi libraries for the four representative strains. For CL-2136, the genome was de novo sequenced using PacBio, and each genome was re-annotated (see Methods). For the NCTC8325-4, JE2, Cowan I, and CL-2136 strains, we designed 2,530, 2,825, 2,764, and 2,580 sgRNAs, respectively, collectively targeting on average 98.6% of annotated genetic features (Supplementary Table 4, Fig. 1D). Genes excluded from the libraries primarily lacked nearby protospacer adjacent motif (PAM) sequences required for CRISPRi targeting. To evaluate sequence divergence and gene content variation among the strains, we quantified the number of genes targeted by sgRNAs with exact sequence matches using a previously established cross-coverage pipeline^16^. The highest overlap was observed between NCTC8325-4 and JE2 (87/93%), consistent with their close genetic relatedness within the same clonal complex. In contrast, overlap declined substantially between JE2 and CL-2136 (62%) and between JE2 and Cowan I (66%), underscoring the strain specificity captured by our CRISPRi libraries across diverse lineages (Supplementary Fig. 1B). Each sgRNA library was synthesized as an oligonucleotide pool containing a universal primer site for amplification of the whole library and a strain-specific primer site for selective amplification (Fig. 1E). The sgRNA cassette within pVL4930 includes flanking Illumina adapter sequences, enabling one-step PCR amplification during library preparation (see Methods). Plasmid pools were first transformed into the restriction-permissive *E. coli* strain IM08B and sequenced to confirm complete sgRNA representation. Subsequently, the libraries were introduced into *S. aureus* strains carrying the *sep::Ptet-dcas9* construct. For each strain, at least 300,000 transformants were collected and pooled to ensure comprehensive library coverage.

### A genome wide-atlas of fitness determinants

CRISPRi-seq was employed to systematically map gene fitness across four *S. aureus* strains. Each strain-specific library was grown in the presence or absence of aTc, in quadruplicate, for 6, 12, and 18 generations. CRISPRi-based fitness estimates were derived from changes in sgRNA abundance following induction (see Methods). Genes with an absolute log_2_FC > 1 (q-value <0.05) were considered to significantly influence bacterial fitness: log_2_FC values < –1 indicate that gene knockdown impaired bacterial fitness (high-fitness genes), whereas log_2_FC values > 1 indicate that knockdown enhanced bacterial fitness (low-fitness genes). CRISPRi induction accounted for most of the variance in sgRNA abundance, whereas biological replicates showed minimal variability. A smaller contribution to variance was associated with the number of replication rounds (Supplementary Fig. 2A). The proportion of fitness genes increased with generation number in each strain, reaching 573 genes for JE2, 581 for Cowan I, 487 for NCTC8325-4, and 560 for CL-2136 after 18 generations (Fig. 2A; Supplementary Fig. 2B and Supplementary Table 5). Gene fitness measurements were broadly consistent across time points, with most fitness-associated genes detectable at 12 generations and remaining identifiable at 18 generations. Correspondingly, sgRNAs targeting high-fitness genes exhibited a progressive decline in log_2_FC values, reflecting enhanced growth inhibition under prolonged CRISPRi-mediated repression (Fig. 2A). An exception was observed for NCTC8325-4 at 12 generations, where a larger number of fitness genes (769) with lower log_2_FC values than in the other strains were detected. Based on these observations, we defined the fitness landscape of each strain as the set of genes whose perturbation conferred a measurable fitness effect after 18 generations. Benchmarking the NCTC8325-4 essentialome against 3 independent transposon sequencing datasets from NCTC-derived strains^28–31^ identified 312 shared high-fitness genes (Supplementary Fig. 2C). Nevertheless, both CRISPRi-seq- and Tn-seq-specific hits were observed, likely reflecting differences in genetic background, growth conditions, screening medium, transposon coverage, and analysis pipelines. For CRISPRi-specific hits, potential dCas9-associated polar effects cannot be excluded. To facilitate exploration of *S. aureus* genomes and essentialomes, we developed a user-friendly web-based genome atlas, AureoBrowse (https://aureobrowse.veeninglab.com). AureoBrowse enables users to search for genes of interest by name or locus tag (Fig. 2F) and includes an interactive track displaying all sgRNAs designed for each strain and their corresponding fitness across experimental conditions. To support comparative analyses, a unique pangenome identifier (panID)^32^ was assigned to each gene.

**Figure 2.**
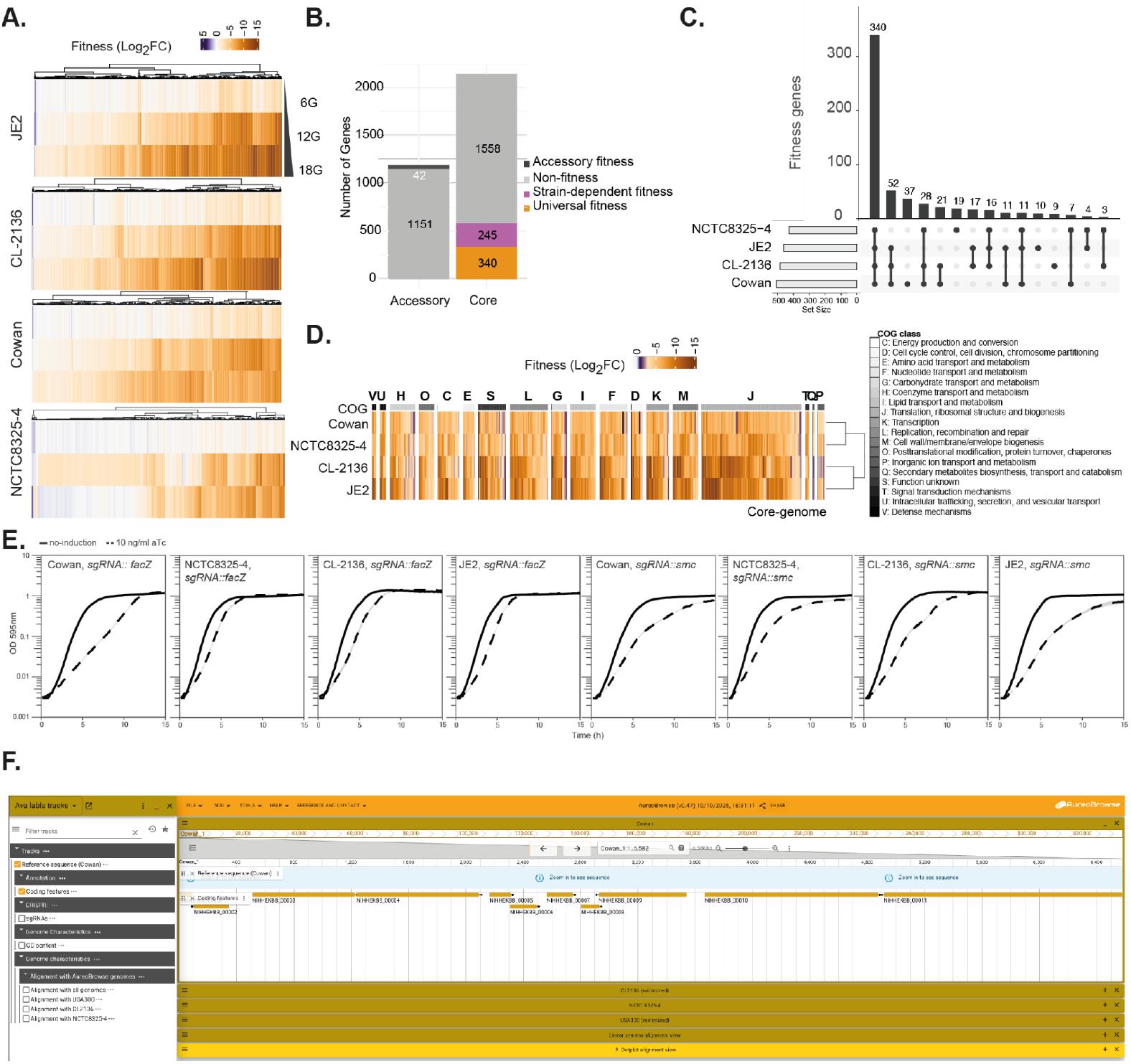
*S. aureus* essentialomes. **A)** Overview of high- and low-fitness genes (|log_2_FC| > 1, *q* < 0.05) across generations for each strain. Negative fitness scores (orange) indicate high-fitness genes, whereas positive scores (purple) indicate low-fitness genes. **B)** Total number of genes in the core and accessory genomes of the four *S. aureus* strains, along with the corresponding numbers of high-fitness genes. Genes were categorized as universal fitness genes (core genes consistently contributing to fitness in all strains), strain-dependent fitness genes (core genes with variable fitness contributions across strains), or accessory fitness genes (accessory genes that strongly affect fitness when present in a strain). **C)** Distribution of fitness genes within the core genome. Horizontal bars represent the number of fitness genes per strain; vertical bars show the number of fitness genes in all four, in subsets, or unique to specific strains. Black dots indicate the strain(s) in which depletion of the genes are causing a fitness defect. **D)** Restricted core essentialome (307 genes; log_2_FC < –1, *q* < 0.05) categorized by COGs functional classes. **E)** Individual targeting of universal (*facZ* and *smc*) fitness genes validates CRISPRi-seq findings. **F)** Screenshot of AureoBrowse interface.

### Strain-specific and core essentialomes

The fitness landscapes were categorized into three classes: (1) accessory fitness genes: accessory genes that strongly affect fitness when present in a strain; (2) strain-dependent fitness genes: core genes with variable fitness contributions across strains; and (3) universal fitness genes: core genes that consistently contribute to fitness in all strains (Fig. 2B and Supplementary Table 5). To establish these classes, the pangenome of the four strains was defined using Roary^33^. Out of 3,328 total genes, 2,144 genes belonged to the core-genome. Across the four strains, we identified 42 accessory fitness genes, highlighting genomic variation and dispensability. Within the core genome, only 73 genes exhibited strict strain-specific essentiality, consistent with overall conservation of core functions^34,35^. However, 245 core genes showed differential fitness impact in pairwise strain comparisons, underscoring the influence of genetic background on essentiality (Fig. 2B, 2C). It is, however, possible that some strain-specific effects may result from dCas9-associated polar effects or limitations in homologue annotation. The universal fitness gene set among the four strains comprised 340 genes. We further refined this by intersecting with a broader *S. aureus* pangenome of 32 strains^32^, defining a narrower core genome of 1,028 genes. This reduced the set to 307 universal fitness genes, which we defined as the core essentialome, representing robust targets for broadly effective antibacterial strategies (Fig. 2D). Within the core essentialome, the three most represented Clusters of Orthologous Groups (COG) categories were translation (COG J; 79 genes), replication, recombination and repair (COG L; 30 genes), and function unknown (COG S; 22 genes), with 22 genes remaining uncharacterized despite manual curation. To validate newly identified universal fitness genes, we targeted *facZ* (cell division and envelope maintenance) and *smc* (chromosome organization and segregation) with the corresponding sgRNA in each strain. Growth assays of the CRISPRi knockdown confirmed their impact on bacterial fitness (Fig. 2E). Their genetic context suggests that the observed phenotypes are unlikely to result from polar effects. Overall, these findings expand the current universal and strain-specific fitness genes landscapes and support context-dependent essentiality.

### CRISPRi-seq chemical genetics to define antibiotic stress signatures

To investigate changes in gene fitness under antibiotic stress, we performed CRISPRi-seq screens in the *S. aureus* Cowan I (MSSA) and JE2 (MRSA) strains using four clinically relevant antibiotics: flucloxacillin (FLUX), vancomycin (VAN), levofloxacin (LEVO), and rifampicin (RIF), representing distinct mechanistic classes (Fig. 3A). Each antibiotic was first tested across a concentration range spanning sub-inhibitory to near-maximal inhibition. Final concentrations were selected based on predicted minimum inhibitory concentrations (MICs) and reduced bacterial growth to 20-40% relative to antibiotic-free controls (Fig. 3B). CRISPRi-seq screens were conducted in the presence or absence of both the inducer (aTc) and each antibiotic. To control for differences in growth kinetics, all cultures were harvested at a uniform optical density corresponding to approximately 12 generations, ensuring an equivalent number of doublings across conditions (Fig. 3A). For flucloxacillin, antibiotic-induced lysis prevented dilution and regrowth to 12 generations; these cultures were therefore harvested after 8 generations (see Methods). Following sgRNA quantification and differential analysis, antibiotic susceptibility determinants were defined as genes exhibiting significant fitness changes relative to the untreated condition (|Δlog_2_FC| > 1, *q* < 0.05), with genes exhibiting Δlog_2_FC < -1 defined as susceptibility genes. Gene fitness profiles were highly reproducible between replicates, and data from 12 generations for LEVO, RIF, and VAN, and 8 generations for FLUX, were used for all subsequent analyses.

**Figure 3.**
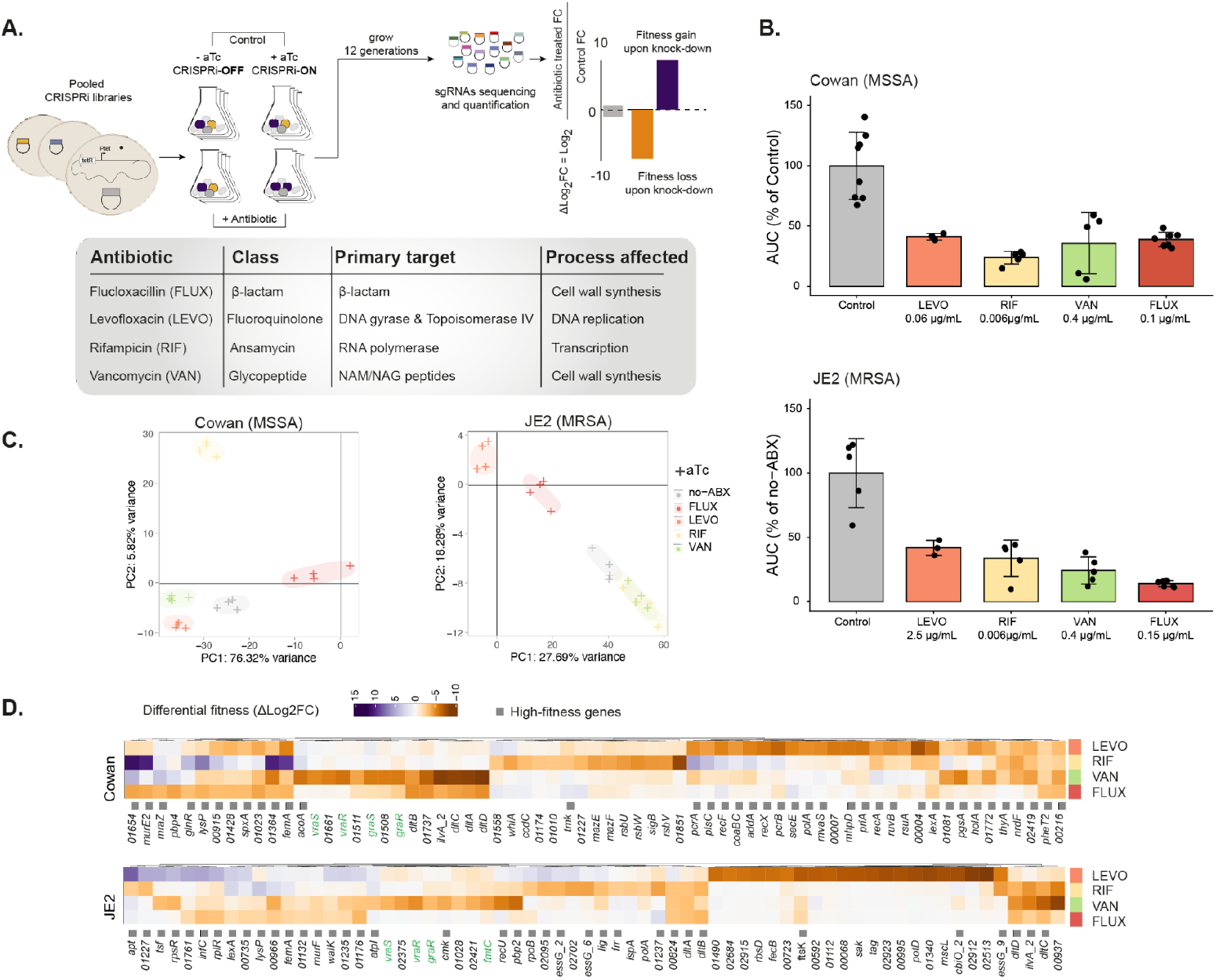
Genome-wide antibiotic genetic signatures. **A)** Schematic of the CRISPRi-seq screening workflow using four clinically relevant antibiotics. **B)** Bacterial survival, expressed as the percentage of the area under the growth curve (AUC) relative to the antibiotic-free control, for Cowan and JE2 strains under control condition and at the indicated concentrations of levofloxacin (LEVO), vancomycin (VAN), rifampicin (RIF), and flucloxacillin (FLUX). **C)** Principal component analysis (PCA) of sgRNA abundances for induced (+aTc) samples showing antibiotic-dependent clustering in both strains. **D)** Differential fitness scores of the 20 top-ranking susceptibility genes (smallest Δlog_2_FC, p < 0.05) in Cowan and JE2 under antibiotic treatment for each of the four antibiotics. Genes causing a relative bacterial fitness loss upon knock-down are shown in orange and genes causing a relative bacterial fitness gain upon knock-down in purple; high-fitness genes identified after 18 generations (no antibiotics) are indicated by grey squares. For both Cowan and JE2 strains, unnamed genes were represented by their Prokka-annotated locus tags (NIHHEKBB and GEBHBEPE respectively), which were omitted from the figure for clarity.

### Antibiotic signatures are genome-wide and related to their mode of action

In both strains, we identified numerous genes exhibiting significant differential fitness (either loss or gain) upon antibiotic treatment (Supplementary Table 6). These findings indicate that antibiotic pressure extends beyond direct drug targets, exerting broad genome-wide effects (Supplementary Fig. 3A). In Cowan I, most differentially affected genes caused fitness loss upon knockdown during levofloxacin (91%) and vancomycin (88%) exposure, whereas a smaller proportion did so in flucloxacillin (42%). In JE2, most genes caused fitness loss upon knockdown in vancomycin (80%) and flucloxacillin (91%), with fewer affected in levofloxacin (31%). In contrast, rifampicin predominantly resulted in fitness gains in both strains, with 72% of affected genes showing increased fitness in Cowan I and 67% in JE2 (Supplementary Table 6). This may reflect the benefit of reduced metabolic burden caused by the global transcriptional inhibition of rifampicin and supports previous observations of strain-specific responses to rifampicin and levofloxacin^36^. Principal component analysis (PCA) of the sgRNA composition across all induced samples revealed that each antibiotic produced distinct susceptibility signatures in both strains, indicative of drug-dependent essentialomes (Fig. 3C). Susceptibility patterns reflected each antibiotic’s mode of action, with 20 top-ranking hits (smallest Δlog_2_FC, p < 0.05) functionally linked to the targeted biological process (Fig. 3D). In Cowan, depletion of genes involved in DNA replication (*priA, pcrA, holA)*, repair (*ruvB, recA*), and recombination (*recX, recF)* caused bacterial fitness defects upon levofloxacin treatment; genes involved in transcription (*sigB)* and translation (*rsbU, rsbV*) were selectively depleted under rifampicin stress; and those associated with cell wall biosynthesis (*mraZ, pbp4*) and envelope homeostasis (*dlt, vraRS, graRS*) were affected by both vancomycin and flucloxacillin (Fig. 3D). In JE2, depletion of the DNA translocase *ftsK* led to a bacterial fitness defect with levofloxacin; *rpoB* and *polA* were key hits with rifampicin; and genes in the *dlt* operon, *murF* (muropeptide), *pbp2* (penicillin binding protein), *vraRS* and *walKR* were associated with susceptibility to vancomycin and flucloxacillin. Several of these well-characterized, expected targets were previously identified in transposon-based screens with vancomycin (green labels, Fig. 3D), corroborating the robustness of our approach. Direct comparison with Tn-seq data for the other antibiotics was not feasible, as previous studies were performed with related compounds from the same classes that differ in their targets and susceptibility mechanisms^7,12,29,30^. Importantly, numerous uncharacterized genes were also highlighted for each class and strain, including high-fitness genes (grey squares, Fig. 3D) that had not been identified in previous chemical-genetic screens.

### Conserved signatures converge in response to drug treatments

Qualitative comparison of genome-wide antibiotic signatures between the Cowan and JE2 core genomes revealed both strain-specific and conserved responses (Supplementary Table 7 and Fig. 4A). While a substantial number of susceptibility genes were shared for vancomycin (78), rifampicin (30), and flucloxacillin (36), overlap was minimal for levofloxacin, with only 12 shared genes (Fig. 4B). This pronounced divergence likely reflects the intrinsic resistance of JE2 to fluoroquinolones ^37^, leading the strain to rely on a distinct set of genes to cope with fluoroquinolone-induced stress. To examine this strain dependency, we generated a clean deletion mutant of *sbcCD*, a putative 3′-5′ exonuclease complex. Growth of the Δ*sbcCD* mutant in the presence of levofloxacin was strongly impaired in JE2, whereas the effect was markedly less pronounced in Cowan (Fig. 4E), thereby confirming the CRISPRi-seq approach. Among the shared susceptibility genes between the two strains, a subset caused synergistic susceptibility across multiple antibiotics (Fig. 4D). A substantial fraction of the antibiotic-shared susceptibility genes (9 of 24) was common to vancomycin and flucloxacillin, consistent with the disruption of cell wall-associated pathways. Depletion of genes involved in cell wall biogenesis, including the *dlt* operon, *ltaS, femA*, and the putative membrane protein SAUPAN002361000, markedly increased susceptibility to both antibiotics, suggesting a role for the latter in cell wall maintenance. In contrast, only two genes, *ftsK* and *pheT2* (tRNA synthetase subunit), led to increased sensitivity to flucloxacillin, levofloxacin, and vancomycin, identifying them as potential core vulnerabilities under general antibiotic stress. We next focused on antibiotic-specific core determinants whose predicted functions align with the respective antibiotic modes of action (Fig. 4E). To further investigate these conserved, antibiotic-specific determinants, we validated the top CRISPRi-seq hits by assessing the growth of the corresponding sgRNA knockdown strains in the presence or absence of the selected antibiotic (FLUX and LEVO). Among the 6 targets selected, 5 were successfully validated using individual sgRNAs, confirming the reliability of the screening (Fig. 5A). Notably, we newly identified *dltB, femB*, involved in peptidoglycan cross-bridging, and *facZ* as conserved susceptibility determinants for FLU. For LEVO, we identified the recombination regulator *recX* and *mvaS*, a key enzyme in the mevalonate pathway, suggesting a potential synergy between LEVO and statins, primary inhibitors of mevalonate synthesis ^38^.

**Figure 4.**
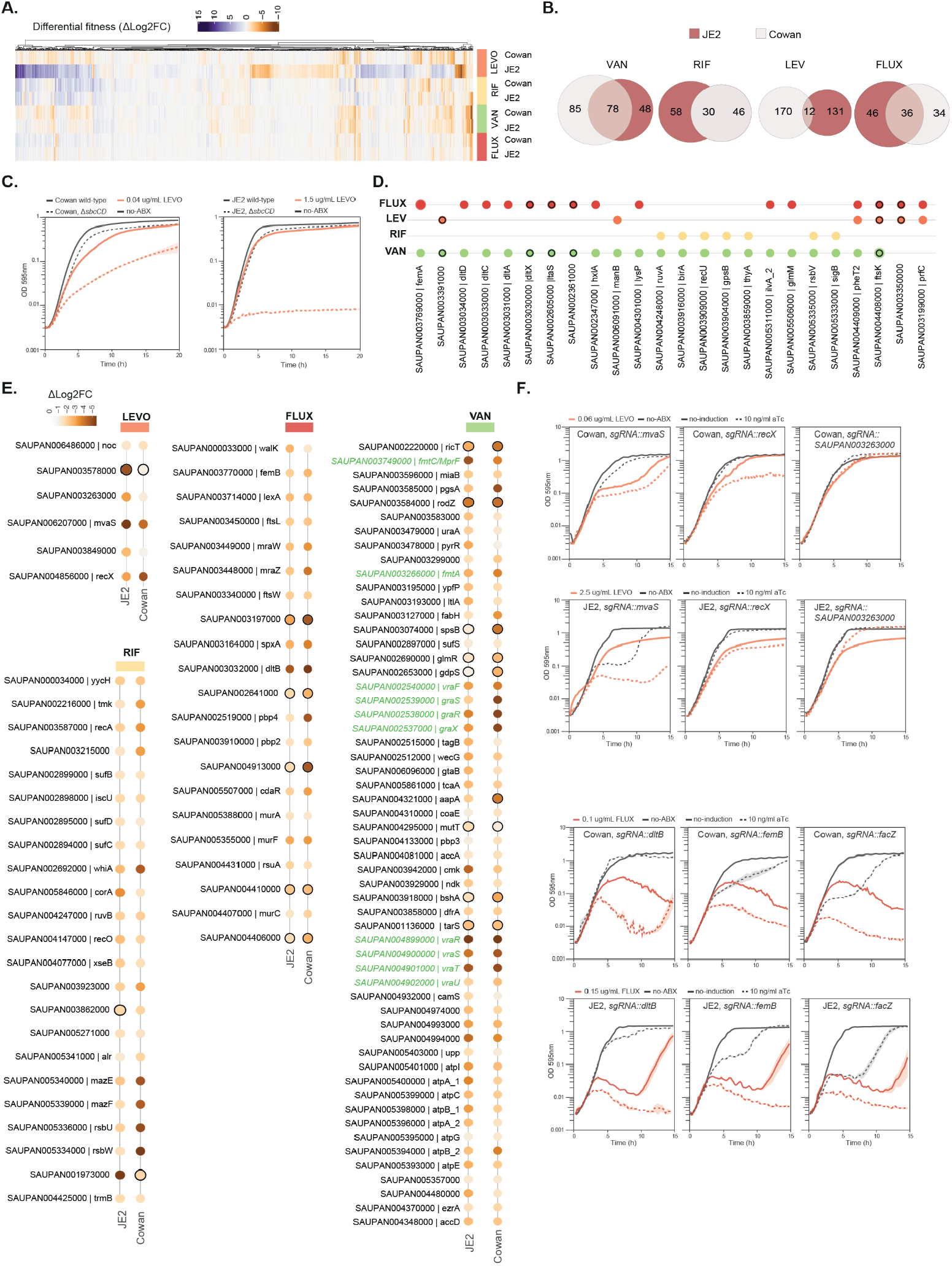
Conserved and strain-specific genetic signatures in antibiotic response. **A)** Differential fitness scores of core genes causing a significant relative bacterial fitness loss upon knock-down (Δlog_2_FC < -1, *q* < 0.05) in at least one strain-antibiotic condition. Genes causing a relative bacterial fitness loss (susceptibility genes) are shown in orange, while those causing a bacterial fitness gain upon knock-down are shown in purple. **B)** Number and overlap of susceptibility genes for each antibiotic in JE2 (red) and Cowan (grey). **C)** Growth curves of JE2 and Cowan wild-type and Δ*sbcCD* confirms strain specificity. **D)** Strain-conserved susceptibility genes (Δlog_2_FC < -1, *q* < 0.05) common to at least two antibiotics. High-fitness genes (18 generations) are outlined with a black circle. **E)** Antibiotic-specific conserved susceptibility genes (Δlog2FC< -1, *q* <0.05). High-fitness genes (18 generations) are outlined with a black circle. **F)** Growth curves confirming the contribution of selected genes to antibiotic susceptibility: *dltB, femB*, and *facZ* (FLUX) and *mvaS, recX*, and *SAUPAN003263000* (LEVO). Gene knockdowns were performed using individual sgRNAs in *S. aureus* JE2 and Cowan strains harboring the *sep::Ptet-dcas9* construct. Growth was monitored under sub-MIC concentrations of the corresponding antibiotics used in the screening.

**Figure 5.**
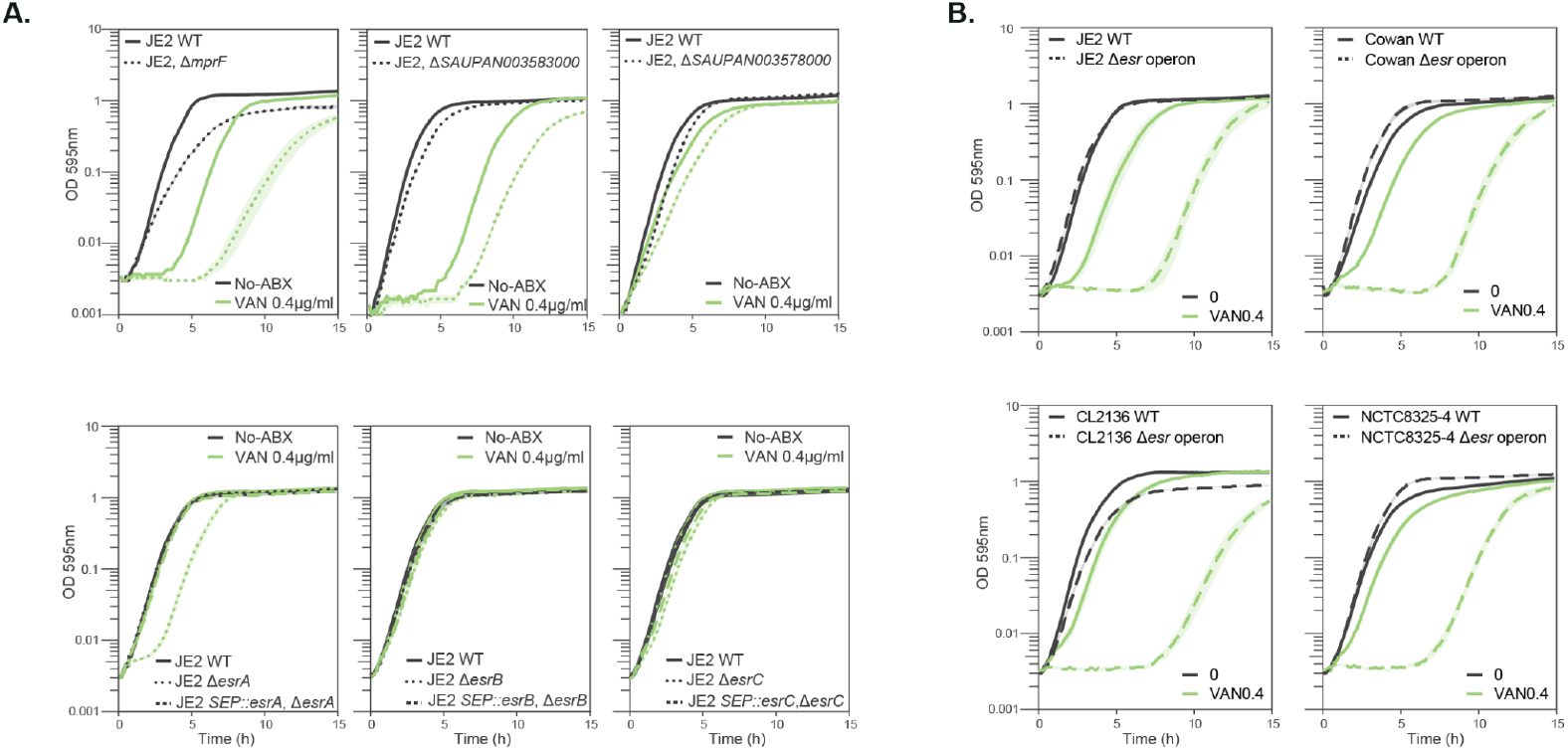
Conserved genetic determinants underlying vancomycin susceptibility. **A)** Growth curves of *S. aureus* JE2 mutants carrying individual deletions of *mrpF*, SAUPAN003583000, SAUPAN003578000, SAUPAN004994000 *(esrA)*, SAUPAN004993000 *(esrB)*, SAUPAN004974000 (*esrC)* and the complementation strains for *esrA, esrB* and *esrC*. **B)** Growth curves of *ΔesrABC* operon deletion mutants in *S. aureus* JE2, Cowan I, CL-2136, and NCTC8325-4 backgrounds confirmed the conserved role of EsrABC in maintaining cell envelope integrity under vancomycin stress.

### Core determinants of vancomycin susceptibility

Vancomycin exhibited the greatest overlap of susceptibility genes, including well-characterized determinants such as *mrpF, fmtA* and the two-component systems *vraRS/graRS* (highlighted in green in Fig. 4E). Several genes identified in this analysis, such as the septation ring regulator *ezrA, ltaS, gpsB, aapA* (Fig. 4D), and SAUPAN004994000 were previously implicated in susceptibility to daptomycin, a lipopeptide antibiotic that also targets the cell envelope^39^. Additionally, previously unrecognized genes emerged as key determinants, including *pbp3* (penicillin-binding protein 3), *rodZ*, a high-fitness gene required for cell envelope integrity and septal organization^40^, and the ATP synthase subunits (*atpD, atpE, atpI*). The latter suggest that inhibition of ATP synthase, as achieved by bedaquiline, might function in synergistic combination with vancomycin^41^. Notably, several genes encoding proteins of unknown function (SAUPAN003299000, SAUPAN004480000, SAUPAN003583000, SAUPAN003578000, SAUPAN004994000, SAUPAN004993000, and SAUPAN004974000) ranked among the strongest vancomycin susceptibility hits (Fig. E). To further investigate these determinants, we generated individual deletions of *mrpF*, SAUPAN003583000 (encoding a GntR-family transcriptional regulator), SAUPAN003578000, SAUPAN004994000, SAUPAN004993000, and SAUPAN004974000 in strain JE2. Growth assays in the presence of vancomycin demonstrated that deletion of any of these genes increased vancomycin susceptibility, further validating our CRISPRi-seq approach (Fig. 5A). The strongest growth defect was observed for SAUPAN004994000, hereafter referred to as EsrA (for envelope stress response factor A). EsrA, together with SAUPAN004993000 (*esrB*) and SAUPAN004974000 (*esrC*), forms an operon co-transcribed from the reverse strand and encoding putative membrane-associated proteins^32^. Deletions of *esrB* and *esrC* produced intermediate phenotypes (Fig. 5A). Complementation of EsrA, B and C with a second, aTc inducible copy expressed at the *sep* locus fully complemented the phenotypes. To assess conservation across genetic backgrounds, the *esrABC* operon was also deleted in the four genetic backgrounds (JE2, Cowan I, CL-2136, and NCTC8325-4). In all strains, deletion of *esrABC* led to markedly increased vancomycin sensitivity, underscoring its conserved and central role in maintaining cell envelope integrity under glycopeptide stress. While further work is required to elucidate the functional roles of these gene products, these findings expand the known repertoire of vancomycin vulnerability genes, implicating multiple and previously unappreciated pathways in maintaining cell envelope integrity.

## Discussion

In this study, we optimized an CRISPRi-seq platform for *S. aureus*, enabling genome-wide investigation of gene fitness across diverse genetic backgrounds. By integrating a tetracycline-inducible, chromosomally encoded *dcas9* construct into a neutral intergenic locus, we achieved tight and tunable repression with minimal toxicity. The system’s responsiveness to doxycycline, a compound with excellent tissue penetration and low host toxicity, further supports its suitability for *in vivo* studies as previously demonstrated for *S. pneumoniae*^*42*^. Through the construction of strain-specific CRISPRi libraries covering nearly all annotated genes, we systematically mapped the fitness landscape in four *S. aureus* isolates representing three major clonal complexes. This analysis defined a conserved set of high-fitness genes shared across strains (core-essentialome), while also revealing extensive variation in gene requirements. Expanding the number of analyzed genomes is expected to further refine, and likely reduce, the size of the core essentialome, emphasizing the plasticity of genetic requirements within the species. For instance, given that the core genome of 1,519 *S. aureus* strains was defined by 162 genes, the core essentialome is likely composed of an equal or smaller number of genes ^43^.

Beyond defining fitness genes, CRISPRi-seq enabled genome-wide chemogenomic profiling under antibiotic stress. Each antibiotic produced a distinct susceptibility signature closely aligned with its mechanism of action: for instance, β-lactam exposure affected cell wall biosynthesis genes such as *mraZ, ftsW*, and *pbp2*, whereas fluoroquinolone treatment perturbed DNA replication and repair pathways including *recX* and *noc*. Importantly, antibiotic-induced stress extended beyond canonical targets, revealing broad cellular adaptations to drug pressure. These findings demonstrate the ability of CRISPRi-seq to define antibiotic stress signatures, offering a scalable approach to dissect the mechanisms of action of uncharacterized compounds. By comparing two genetically divergent strains, the MRSA strains JE2 and the MSSA strain Cowan I, we identified both shared and lineage-specific antibiotic responses. The extent of divergence observed for flucloxacillin, and levofloxacin signatures likely reflects strain-specific resistance determinants such as the presence of *mecA* and mutations in topoisomerases^25^. For instance, the exonuclease complex *sbcCD* appeared critical for maintaining genome stability under fluoroquinolone stress in the resistant JE2 background, possibly reflecting a compensatory role linked to topoisomerase mutations^25^. Conversely, depletion of a single operon, *pheT2-ftsK*, sensitized both strains to three antibiotics (flucloxacillin, levofloxacin, and vancomycin) indicating a potential role as a general sensitizing factor to antibiotic stress. However, we did not identify a specific mechanism conferring intrinsic resistance to multiple drugs, such as those described in *Mycobacterium tuberculosis*^*44*^. Validation of individual hits largely confirmed the pooled screening results; however, discrepancies may reflect differences between pooled and single-clone assays. Sub-inhibitory concentrations of antibiotics can induce the formation of small-colony variants^45^, which may confer relative fitness advantages detectable only in pooled populations, thereby explaining such variations.

Vancomycin produced the most extensive set of susceptibility hits. Among these, we identified a previously uncharacterized operon, *esrABC*, encoding three predicted membrane proteins. Individual deletions of *esrA, esrB*, and *esrC* revealed that *esrA* contributes most strongly to susceptibility, while deletions of *esrB* and *esrC* caused intermediate defects. The operon is conserved across *S. aureus* lineages, and has been implicated in susceptibility to other cell-envelope-targeting antibiotics, including daptomycin and dalbavancin^7,17^. Although the biological function of EsrABC remains to be elucidated, its co-transcription and predicted topology suggest that it forms a membrane-associated complex involved in envelope stress adaptation. It would be of particular interest to determine whether deletion of this complex in VISA or VRSA strains resensitizes them to vancomycin. Together, this work provides a comprehensive atlas of *S. aureus* fitness and antibiotic responses across multiple strains. Beyond revealing strain-specific dependencies, our data highlight conserved targets that shed light on antibiotic modes of action and resistance. Future efforts to characterize newly identified systems, such as EsrABC, and to leverage these insights through combination therapies may offer new avenues to combat antibiotic-resistant *S. aureus*.

## Materials and methods

### Bacterial strains, plasmids and culture conditions

Bacterial strains and plasmids used in this study are listed in Supplementary Table 1. *S. aureus* strains were cultured in tryptic soy broth (TSB) or agar (TSA) and incubated at 37°C and 200 rpm. *Escherichia coli* strains were cultured in lysogeny broth (LB) at 37°C and 200 rpm. When required, growth media were supplemented with trimethoprim (TMP) (10 μg mL-1), chloramphenicol (CHL) (optimized at 20 μg mL-1), anhydrotetracycline hydrochloride (ATC) (20 ng mL-1), erythromycin (Ery) (5 μg mL-1), kanamycin (KAN) (50 μg mL-1), Isopropyl β-Dthiogalactopyranoside (IPTG) (0.5 mM), or X-gal (150 μg mL-1).

### Strain constructions

Strains were constructed using either pMAD- or pIMAY-based allelic exchange systems as previously described^46,47^, or by recombineering. Detailed information on strain and plasmid construction is provided in Supplementary Table 3. The ptet-dcas9 construct and the second copies of selected genes (complementation strains) were inserted into the newly designated *sep* locus, a conserved intergenic region flanked by SAUPAN006480000 and SAUPAN006481000. For recombineering, electrocompetent *S. aureus* cells carrying the recombineering plasmid pPM260 (Supplementary Table 1) were prepared as follows. Overnight cultures grown in TSB supplemented with erythromycin were diluted 1:100 into fresh TSB containing 0.5 M sorbitol (TSBS) and erythromycin. Cultures were incubated at 37 °C until reaching an optical density at 600 nm (OD_600_) of 0.3, at which point IPTG (0.5 mM) was added to induce expression of the recombineering machinery. Growth was continued until an OD_600_ of 0.8 was reached. Cells were harvested by centrifugation at 4000 rpm for 10 min at room temperature, washed twice with an equal volume of sterile 0.5 M sucrose solution, and resuspended in

0.5 M sucrose to a final volume of 400 µL. Competent cells (200 µL aliquots) were used immediately or stored at –80 °C. Linear DNA fragments containing two ∼500 bp homology arms flanking the construct of interest were generated by Golden Gate assembly, amplified by PCR, and dialyzed (primers are listed in Supplementary Table 2). The resulting fragments were electroporated into *S. aureus* cells carrying pPM260 using 2 mm cuvettes and the following parameters: 2.9 kV, 100 Ω, 25 µF. Homologous recombination allowed chromosomal integration at the *sep* or native target locus. After electroporation, cells were recovered in TSBS for 3 h at 37 °C and plated on TSA containing the appropriate antibiotic for selection of integrants. After overnight incubation, colony PCR was performed to identify positive clones. Confirmed clones were cultured overnight under the same selective conditions, with IPTG (0.5 mM) added when necessary to promote loss of the recombineering plasmid. Cultures were then streaked onto selective plates to isolate individual colonies. Loss of the recombineering plasmid was verified by replica plating onto TSA containing either the cassette-selective antibiotic or erythromycin and assessing antibiotic sensitivity. Colonies exhibiting the expected sensitivity profile were confirmed by Sanger sequencing to ensure correct chromosomal integration. For preparation of electrocompetent cells without the recombineering system, the IPTG induction step was omitted, and cells were harvested at OD_600_ = 0.5. Subsequently, cells were resuspended and harvested sequentially in 1/10th and 1/25th of the original culture volume in 0.5M sucrose solution. Finally, the pellet was resuspended in 0.5M sucrose solution to a final volume of 400 μL. Competent cells were aliquoted (200 μL per tube) and either used immediately or stored at –80 °C.

### Construction of single sgRNAs plasmids

The 20 bp target-specific region of each sgRNA was inserted into the plasmid pVL4930 (Supplementary Table 3) by Golden Gate assembly, following a previously described procedure^42^. Briefly, complementary oligonucleotides for each sgRNA (Table SX) were mixed in TEN buffer (10 mM Tris, 1 mM EDTA, 100 mM NaCl, pH 8), heated to 95 °C for 5 min, and allowed to gradually cool to room temperature to enable annealing. The recipient plasmid pVL4930 was digested with BsmBI to excise the mCherry fragment and purified by gel extraction. Annealed oligos were then ligated into the linearized vector using T4 DNA ligase and transformed into *E. coli* IM08B. Plasmid constructs were confirmed by Sanger sequencing with primer OVL2153.

### Design and construction of strain-specific sgRNA libraries

The genome of the clinical isolate CL-2136 was sequenced using PacBio technology, and all the genetic features were annotated using Prokka. The publicly available genomes of the other three strains (accession numbers NC_007795, GCF_00013425 and GCF_016916815.1) were also re-annotated with the same pipeline. Based on the newly annotated features, we applied our recently published pipeline for de novo sgRNA design ^16^. The libraries were designed to include a single optimal sgRNA per target, minimizing off-target effects and accounting for the distance from the transcription start site. The pooled libraries were synthesized by Twist Bioscience. For each strain, sgRNAs were amplified from the Twist-synthesized library using strain-specific primers (Supplementary Table 2). Amplified fragments (∼92 bp) were purified with the Monarch PCR & DNA Cleanup Kit (New England Biolabs) according to the manufacturer’s protocol. The vector pVL4930 was linearized by BsmBI digestion within the mCherry locus and used as a backbone for library assembly. Golden Gate cloning was performed using BsmBI and T4 DNA ligase (Vazyme) with a vector:insert ratio of 1:25, cycling between 37 °C and 16 °C for 70 cycles, followed by heat inactivation at 80°C. Reaction mixtures were dialyzed against sterile water prior to transformation. Assembled plasmid libraries were electroporated into *E. coli* Stbl3, recovered in LB, and plated on selective media. Successful replacement of mCherry was indicated by white colony color, with cloning efficiencies typically exceeding 50.000 colonies per reaction. Library diversity was assessed by colony PCR and confirmed by Sanger sequencing using primer OVL2153. For long-term storage, bacterial colonies were pooled, mixed 1:4 with 80% glycerol, and stored at -80 °C. Library complexity was validated by Illumina sequencing, ensuring comprehensive representation of sgRNAs prior to transformation into *S. aureus* strains. Plasmid libraries isolated from *E. coli* were introduced into *S. aureus* JE2, NCTC8325-4, Cowan I, and CL-2136 strains carrying the *sep::Ptet-dcas9* construct by electroporation. Multiple parallel transformations were performed and plated on tryptic soy agar (TSA) supplemented with 10 µg/mL chloramphenicol. Approximately 3 × 105 colonies were harvested using a cell scraper, pooled, and resuspended in ∼50 mL TSB containing chloramphenicol (10 µg/mL). The combined libraries were subsequently diluted (500 µL into 2 L Erlenmeyer flasks containing 500 mL TSB with 10 µg/mL chloramphenicol), cultured overnight at 37 °C with agitation, and preserved as glycerol stocks for subsequent experiments.

### CRISPRi-seq screens

Cultures were diluted to an initial OD_600_ of 0.01 in fresh TSB containing chloramphenicol (10 µg/mL) and grown to OD_600_ 0.3 to generate starter cultures. For each screening condition, 50 mL of TSB supplemented with chloramphenicol (10 µg/mL), with or without inducer (10 ng/mL anhydrotetracycline or doxycycline), was inoculated at an initial OD_600_ of 0.01. Cultures were incubated in 200 mL flasks at 37 °C with shaking (200 rpm) and serially propagated for 6, 12, and 18 generations by successive 1:100 back-dilution into fresh medium at mid-exponential phase (OD_600_ ≈ 0.6). At each generation point, cells were harvested by centrifugation (4000 rpm, 10 min, 4 °C), resuspended in 1 mL of TSB containing 20% glycerol, and stored at -80 °C for plasmid extraction and downstream analyses. For each time point and condition, four biological replicates were performed.

#### Screenings with levofloxacin, rifampicin and vancomycin

The frozen library stock was diluted 1:1000 in 50 mL of medium containing chloramphenicol and cultured overnight. The overnight culture was then diluted to an OD600 ≈ 0.01 in 30 mL and grown as a preculture to an OD600 ≈ 0.3, corresponding to exponential growth. Cells were further diluted to an OD600 ≈ 0.01 and distributed into four replicates of 30 mL of cultures for each of the following growth conditions: (a) with antibiotics with aTc, (b) with antibiotics without aTc, (c) without antibiotics with aTc, and (d) without antibiotics without aTc. The antibiotics levofloxacin, rifampicin, and vancomycin were used at sub-MIC concentrations previously determined for the strains Cowan and JE2. Cultures were grown to an OD600 ≈ 0.6 (approximately 6 generations), at which point they were used to inoculate fresh cultures under the same conditions at an initial OD600 ≈ 0.01. The remaining culture was stored. Newly inoculated cultures were grown again to an OD600 ≈ 0.6 (approximately 12 generations) before being stored.

#### Screenings with flucloxacillin

Overnight cultures and precultures were prepared as described for the other antibiotics. Cells were diluted to an initial OD600 ≈ 0.003 and distributed into four replicates of 30 mL of culture for each of the same growth conditions (a), (b), (c), and (d). Flucloxacillin was used at sub-MIC concentrations determined for Cowan and JE2. Cultures were grown to an OD600 ≈ 0.4 (approximately 8 generations), before being stored.

### Library preparation and Illumina sequencing

Samples from screening cultures were centrifuged into pellets (4000 x g, 10 min, 4°C) and stored in TSB with 20% glycerol at -80°C. For bacterial lysis, the frozen cells were resuspended in 500 μL of Tris-EDTA buffer. 0.8 mg/mL of lysozyme and 0.02 mg/mL of lysostaphin were added and the samples were incubated for 1h at 37°C. For plasmid-isolation, a miniprep kit for *S. aureus* (PureYield Plasmid Miniprepkit System, Promega) was used. DNA was stored at -20°C. sgRNAs from the minipreps were amplified using Nextera DNA Index primers, each containing an Illumina barcode. Different primer pairs were used for the various samples from different growth conditions, allowing accurate assignment of sequencing reads to the respective condition. The amplification of sgRNAs was performed through 25 cycles of PCR. The PCR-products were purified using an agarose gel extraction kit (PCR clean-up Gel extraction, Macherey-Nagel) and quantified by Qubit DNA quantification. The samples were then normalized in RSB, pooled and sent to the UNIL Genomic Technologies Facility for sequencing. The pooled samples were sequenced on the NovaSeq platform using 150 bp reads.

### CRISPRi-seq analysis

The number of reads for each sgRNA was quantified using the software 2FAST2Q^48^ using default parameters. The extracted counts were further analyzed using the R package DESeq2 to determine enrichment or depletion^49^. Tests were made against an absolute log2FC of 1, with a significance threshold of α = 0.05 applied.

### Pangenome analysis

To determine the pangenome of the four strains, Prokka-annotated fasta files were used as input for Roary using default parameters^33^. Pangenome Identifier (PAN ID) was added to each homologue and used as comparative identifier. To compare the core-essentialome to the pangenome of the 32 strains, we used the homologues table from AureoWiki^32^.

### MIC determination

Wild-type Cowan and JE2 strains were cultured overnight in 10 mL volumes without exposure to antibiotics. The overnight cultures were diluted to an OD600 ≈ 0.01 in 30 mL of volume and grown as a preculture to an OD600 ≈ 0.3 corresponding to exponential growth. The cells were then diluted into

30 mL cultures containing different concentrations of the antibiotics flucloxacillin, rifampicin, vancomycin or levofloxacin. The concentrations tested range from 0.02 to 0.2 μg/mL for flucloxacillin, 0.003 to 0.1 μg/mL for rifampicin, 0.2 to 0.5 μg/mL for vancomycin and 0.06 to 3.2 µg/mL for levofloxacin. Initial inoculation densities were set to an OD600 ≈ 0.01 for rifampicin and vancomycin and 0.003 for flucloxacillin. Bacterial growth was measured every hour over a period of 5 to 6 hours. The growth curves for the different antibiotic concentrations were generated, and bacterial survival was assessed by the area under the curve (AUC). The concentration was defined as a concentration below the MIC that causes a distinct growth defect compared to cells grown without antibiotics. A distinct growth defect was defined as a reduction of bacterial survival to 20 to 40% of the levels observed in antibiotic-free medium. For Cowan, sub-MICs were determined as 0.1, 0.4, 0.006, and 0.06 µg/mL for flucloxacillin, rifampicin, vancomycin, and levofloxacin, respectively, and for JE2 as 0.15, 0.4, 0.006, and 2.5 µg/mL, respectively.

### Microtiter plate-based growth assay

Growth assays were performed by first growing the frozen cells in TSB medium at 37°C until mid-exponential growth phase (OD600 = 0.2-0.3). Cultures were then diluted 100-fold into TSB medium and with or without the addition of anydrotetracycline (10 ng/ml) when needed, and antibacterial compounds as indicated in the main text and/or figures and/or figure legends. 150 µL of bacterial culture was then transferred in triplicate (technical replicates) into 96-well plates. Growth was monitored by measuring the OD600 every 10 min for 20 hr at 37°C without CO2 in a TECAN Infinite 662 F200 Pro. OD600 values were normalized so that the lowest value measured during the first hour of growth was 0.004, the initial OD600 value of the inoculum. Each growth assay was performed in triplicate (biological replicate) and the mean value plotted, with the SEM (Standard Error of the Mean) as shading using BactExtract.

### AureoBrowse

AureoBrowse (https://aureobrowse.veeninglab.com) is based on JBrowse ^50^. Features were divided over three annotation tracks: (i) reference sequence, (ii) coding features, and (iii) designed sgRNAs. GC content is calculated via the NucContent plugin (available at https://github.com/jjrozewicki/jbrowse2-plugin-nuccontent).

## Acknowledgements

Work in the Veening lab is supported by the Swiss National Science Foundation (SNSF) (project grants 310030_192517, 310030_200792, 310030_204343 and ‘AntiResist’ 51NF40_180541). The funders had no role in study design, data collection and analysis, decision to publish, or preparation of the manuscript. We thank Geraldine Menoud, Lia Da Rold, Jazmine Mote, Kerstin Strengen and Sandro Jakonia for their contribution in determining sub-MIC antibiotic concentrations and generating mutant strains. We thank Andrew Varble for sharing the recombeeniring plasmids and protocol. We thank the Genome Technologies Facility (GTF) department in Lausanne for the sequencing.

## Authors Contributions

M.V.M wrote the paper with input from all authors. M.V.M, J.B, A.S, L.M. performed the experiments. M.V.M, J.B. and J.W.V designed, analysed and interpreted the data.

