## Supplementary Figure 1 for "Strain-resolved CRISPRi-Seq reveals conserved antibiotic vulnerabilities in *Staphylococcus aureus*"

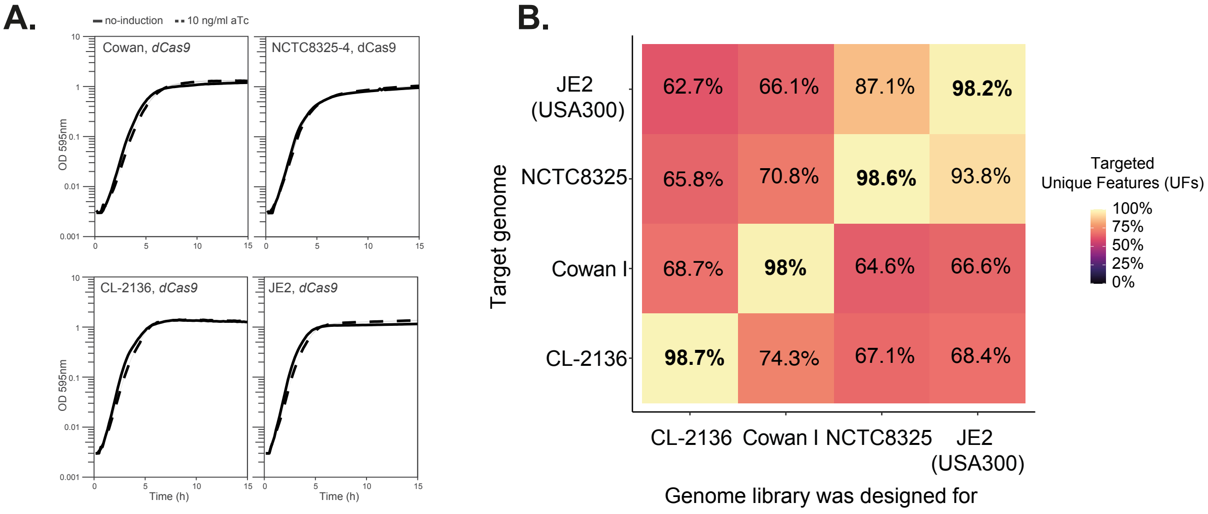


**Supplementary Figure 1.** A) Growth curves of NCTC8325-4, JE2, Cowan I, and CL-2136 strains harbouring sep::Ptet-dcas9. B) Genome cross-coverage of each CRISPRi library. The percentage of sgRNAs uniquely targeting exact genomic sequences (no mismatches; unique targets, UFs) is shown for each pairwise strain comparison.
