## Supplementary Figure 2 for "Strain-resolved CRISPRi-Seq reveals conserved antibiotic vulnerabilities in *Staphylococcus aureus*"

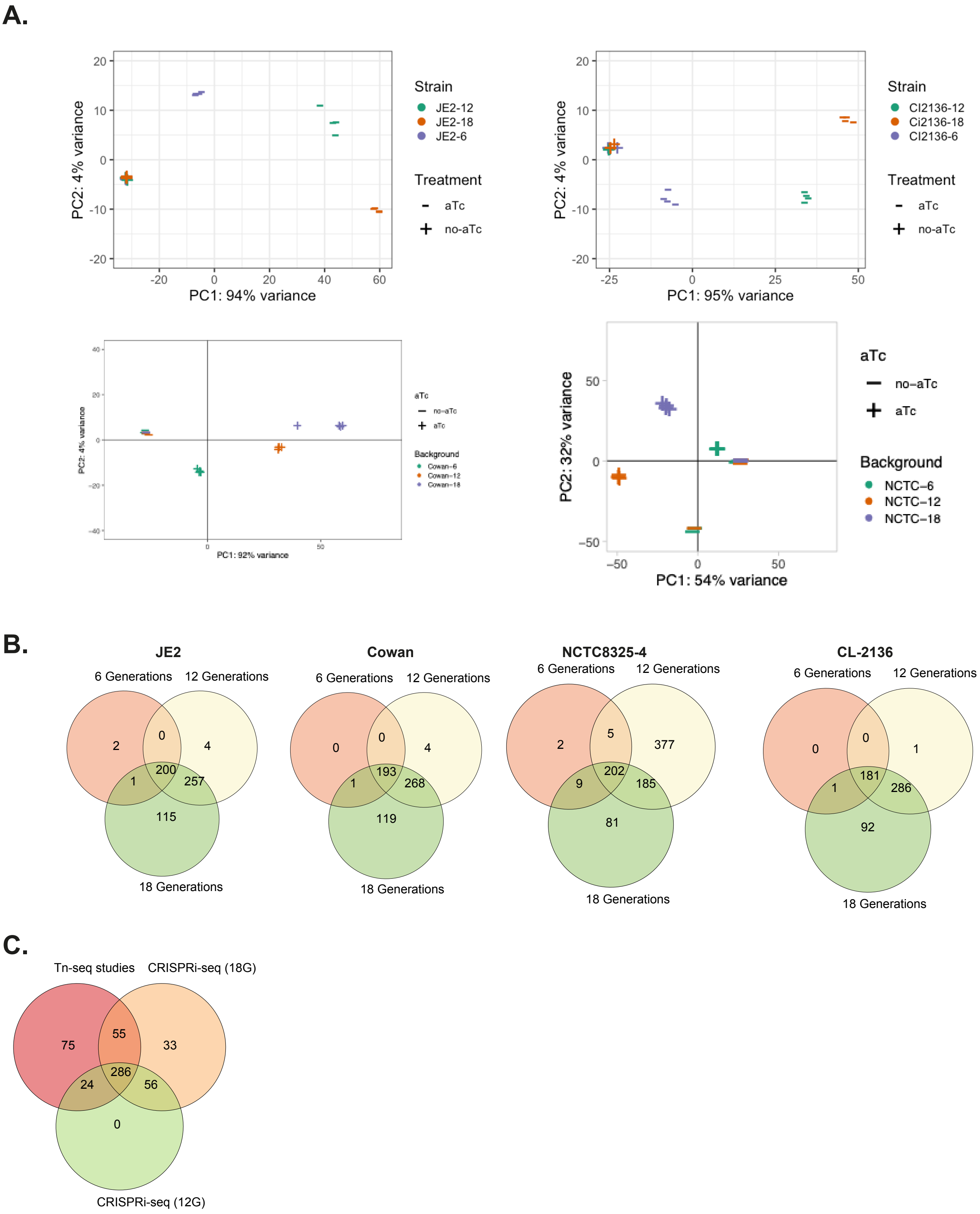


**Supplementary Figure 2.** A) Principal Component Analysis of the sgRNA counts at 6, 12, 18 generations for each strain. B) Venn diagram showing the overlap of fitness genes at 6, 12, 18 generations for each strain C) Venn diagram showing the comparison of genes with significant fitness defect by CRISPRi-seq in NCTC8325-4 (12 and 18 generations) and essentialomes as defined by Tn-seq-based screens of related strains25–28.
