## Supplementary Figure 3 for "Strain-resolved CRISPRi-Seq reveals conserved antibiotic vulnerabilities in *Staphylococcus aureus*"

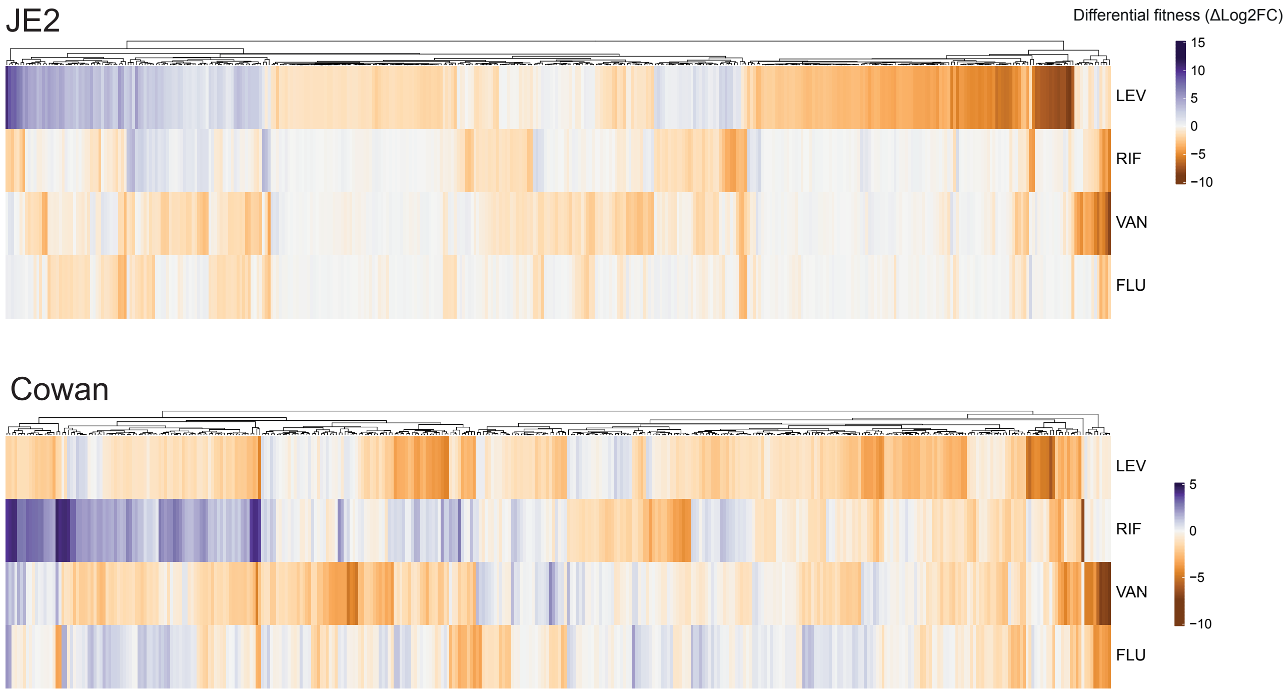


**Supplementary Figure 3.** Genome-wide profiling of antibiotic stress signatures for JE2 and Cowan. Fitness scores of sgRNA targets causing a significant bacterial fitness loss (Δlog2FC< -1, q <0.05) upon knock-down in at least one antibiotic.
